# A hybrid nitrogenase with regulatory elasticity in *Azotobacter vinelandii*

**DOI:** 10.1101/2023.06.02.543473

**Authors:** Alex J. Rivier, Kevin S. Myers, Amanda K. Garcia, Morgan S. Sobol, Betül Kaçar

## Abstract

Biological nitrogen fixation, the microbial reduction of atmospheric nitrogen to bioavailable ammonia, represents both a major limitation on biological productivity and a highly desirable engineering target for synthetic biology. However, engineering of nitrogen fixation requires an integrated understanding of how the gene regulatory dynamics of host diazotrophs restrict the available sequence-function space of its central catalytic metalloenzyme, nitrogenase. Here, we interrogate this relationship by analyzing the transcriptome of *Azotobacter vinelandii* engineered with a phylogenetically inferred, ancestral nitrogenase protein variant. The engineered strain exhibits reduced cellular nitrogenase activity but recovers wild-type growth rates following an extended lag period. We find that expression of genes within the immediate nitrogen fixation network is resilient to nitrogenase sequence-level perturbations. Rather, physiological compatibility with the ancestral nitrogenase variant is restored by reducing trace metal and electron resource allocation to nitrogenase. Our results spotlight cellular processes adjacent to nitrogen fixation as productive engineering targets to improve compatibility between remodeled nitrogenase proteins and engineered host diazotrophs.

**IMPORTANCE:** *Azotobacter vinelandii* is a key model bacterium for the study of biological nitrogen fixation, an important metabolic process catalyzed by nitrogenase enzymes. Here, we demonstrate that compatibilities between engineered *A. vinelandii* strains and remodeled nitrogenase variants can be modulated at the regulatory level. Engineered cells respond by adjusting expression of proteins involved in cellular processes adjacent to nitrogen fixation, rather than that of nitrogenase proteins themselves. These insights can inform future strategies to transfer nitrogenase variants to non-native hosts.

## INTRODUCTION

Nitrogen cycling impacts ecosystems across the globe and is vitally important for sustained biological activity. The largest reservoir of nitrogen is highly inert, atmospheric N_2_ that is unavailable to most organisms. Nature has invented a single molecular mechanism to reduce, or “fix”, N_2_ gas to bioavailable NH_3_ and overcome nitrogen limitations on biological productivity via the family of nitrogenase enzymes hosted solely by certain bacteria and archaea (1, 2). Nevertheless, approximately half of global fixed nitrogen today is generated by anthropogenic means to meet the demands of a rapidly expanding human population (3). Whereas nitrogenases catalyze nitrogen fixation at ambient conditions, the Haber-Bosch process, which generates the bulk of anthropogenic fixed nitrogen, requires high temperatures and pressures and is both energetically and environmentally costly (4). Thus, strategies to both improve biological nitrogen fixation activity and to distribute the enzymatic machinery to non-diazotrophic hosts (e.g., cereal crops) are highly desirable bioengineering goals (5, 6).

A critical component of nitrogen fixation in natural diazotrophs is its genetic regulatory architecture that is required to coordinate the expression of nitrogenase enzymes in response to dynamic environmental conditions and physiological states (7–9). Regulatory precision is necessary due to the high metabolic cost of nitrogen fixation as well as the complex nature of its supporting protein network. For example, the aerobic, diazotrophic model gammaproteobacterium *Azotobacter vinelandii* (*A. vinelandii*) has more than 50 proteins that support three nitrogenase isozyme systems, with molecular functions including nitrogenase regulation and assembly, electron transport, and cofactor synthesis (8, 10, 11). The oxygen sensitivity of the nitrogenase metalloenzyme demands that aerobic microbial hosts (desirable models for nitrogen fixation transfer to crop plants such as cereals) utilize particular strategies to ensure metabolic compatibility, including temporal/spatial regulation of nitrogen fixation activity or protection via heightened respiratory rates (9, 12). These features make regulatory optimization challenging, particularly with heterologous expression of nitrogen fixation genes. Under diazotrophic conditions (i.e., lacking an exogenous reduced nitrogen source), nitrogenases are highly expressed (e.g., comprising up to 10% of total protein in *A. vinelandii* (13)), and, counterintuitively, overexpression typically decreases total nitrogen fixation activity (14), perhaps due to misfolding of excess proteins (15). Nitrogen fixation evidently relies on a fine-tuned protein network stoichiometry that is well-optimized in native hosts but is often incompatible with other species (16). Attempts at rational and/or combinatorial reengineering of nitrogen fixation regulatory schemes have revealed that their underlying principles are not fully understood.

Relative to regulation, sequence-level engineering of nitrogenases and their associated proteins for improved outcomes has received little attention. Prior mutagenesis studies have generally aimed to generate key insights into nitrogenase biochemical mechanism via disruption of activity rather than to increase activity through a broader exploration of sequence space (1, 17). Only recently have the functional consequences of nitrogenase sequence variation been deeply interrogated, for example, through random mutagenesis (18), extant ortholog libraries (19), and resurrection of phylogenetically inferred, ancestral proteins (20). These sequence-level studies have the potential to identify variants with improved catalytic activities, stabilities, or interactions with associated proteins, as well as compromises of two or more of these features (e.g., reduced activity but highly improved stability). Nevertheless, due to the many genetic requirements for nitrogenase activity, functional insights from these studies are unlikely to be divorced from their downstream consequences on the surrounding cellular network in both native and heterologous hosts. Sequence-level optimization might be hampered by protein network incompatibilities and/or a counterproductive regulatory response (21). Thus, successful nitrogen fixation bioengineering prospects rely upon a concerted understanding of both nitrogenase sequence and regulatory space.

In this study, we probe the regulatory response of *A. vinelandii* to a synthetic, ancestral variant of the nitrogenase catalytic protein NifD, previously resurrected from within the direct *A. vinelandii* evolutionary lineage (20). The establishment of the *A. vinelandii* genetic system, which has detailed the nitrogenase regulon structure and expression mechanisms, and the detailed physiological information available from decades of work, makes *A. vinelandii* an ideal model organism for the transformation of synthetic nitrogenase genes and downstream transcriptional characterization (1, 8, 10, 22, 23). The engineered *A. vinelandii* strain is capable of diazotrophy but exhibits perturbed growth and nitrogenase substrate reduction rates. By analyzing the transcriptome of the engineered strain, we identify the specific genetic components most likely to contribute to the phenotype. We find that transcription levels within the immediate nitrogen fixation network, including nitrogenase and nitrogenase-related gene clusters, are highly robust to mutations in a core, catalytic component of nitrogen fixation. Rather, the regulatory response is enriched for genes external to this immediate network that indirectly impact nitrogen fixation by modulating electron flux, trace metal transport, motility, stress response, and central metabolism. Our results shift focus to these ancillary cellular functions as potential engineering targets to improve compatibility of remodeled nitrogenase proteins in diazotrophic hosts.

## RESULTS & DISCUSSION

### An ancestral nitrogenase protein variant results in defects to *A. vinelandii* nitrogen fixation ability

To interrogate the regulatory consequences of sequence-level changes in an engineered nitrogenase gene, we selected a previously constructed *A. vinelandii* strain, “ancNif”, in which one of the wild-type (WT) nitrogenase catalytic genes, *nifD* (encoding the NifD protein subunit), was replaced with a phylogenetically inferred, ancestral variant (20, 24, 25). The ancestral NifD protein sequence was reconstructed from a phylogenetic node within the extant *A. vinelandii* nitrogenase evolutionary lineage and bears ∼85% protein sequence identity to WT NifD (**SI File 1 – Fig. S1, Fig. S2**). Unlike random mutagenesis or replacement with extant orthologs from divergent clades, our strategy introduces sequence variation derived from the evolutionary relationships across extant nitrogenases within a proteobacteria clade and thus mode likely to retain genomic compatibility (20).

NifD, together with NifH and NifK subunit proteins, comprise the molybdenum- (Mo-) nitrogenase complex. Whereas *A. vinelandii* also possesses genes encoding vanadium- (V-) and iron- (Fe-)nitrogenase isozymes, which differ in their metal dependencies and expression conditions (26), only *nifD* has been replaced with an ancestral variant in the ancNif strain. Therefore, the Mo-nitrogenase in ancNif is a hybrid enzyme complex of ancestral and WT subunits (**Fig. 1A**). The NifD protein subunit binds the iron-molybdenum metallocluster (FeMo-co) that serves as the site for nitrogenase substrate reduction and together with NifK comprises the heterotetrameric catalytic component of nitrogenase. Electrons are delivered to the NifD active site via transient interactions between NifDK and the homodimeric nitrogenase reductase component NifH.

**Fig. 1.**
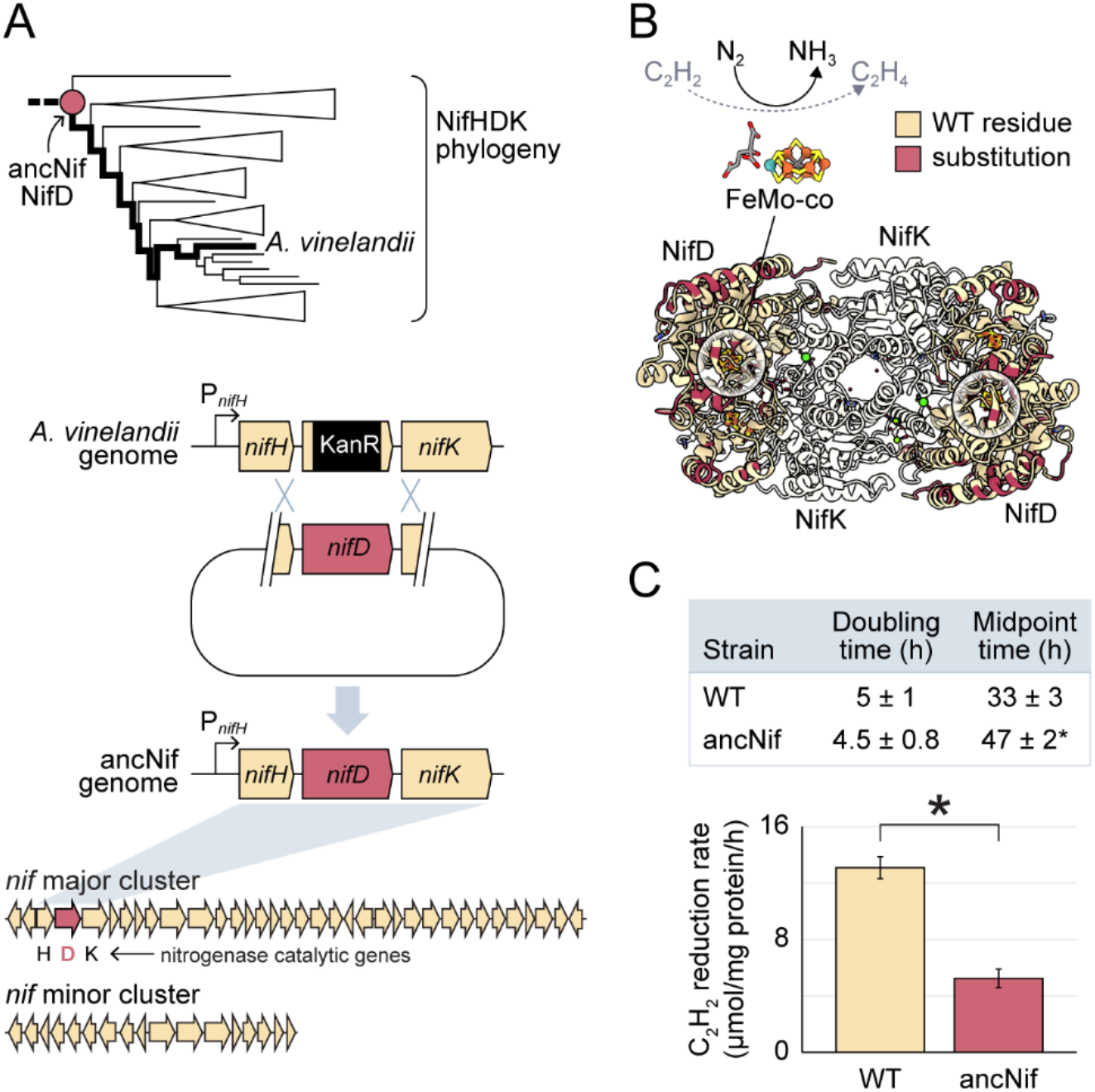
Construction and physiology of *A. vinelandii* strain ancNif harboring an ancestral nitrogenase NifD protein subunit. (A) The protein sequence of ancestral NifD was inferred from a NifHDK protein phylogeny (the targeted ancestral NifD clade, which includes the *A. vinelandii* lineage, bold, is shown). The ancestral *nifD* gene (pink) was integrated into the *A. vinelandii* genome by homologous recombination, replacing a kanamycin resistance marker (KanR) previously incorporated to knock out WT *nifD* (see **Materials and Methods**). The engineered ancestral gene is the only genetic perturbation within the broader *nif* major and minor clusters. (B) ColabFold-predicted structure of the hybrid nitrogenase catalytic tetramer, NifDK, in ancNif. NifD subunits are colored tan, with residues within the ancestral NifD that are substituted relative to WT highlighted pink. NifK is shown transparent. FeMo-co serves as both the site of N_2_ reduction to NH_3_, as well as reduction of the alternative substrate C_2_H_2_ to C_2_H_4_ (dotted arrow). (C) Tabulated growth parameters of ancNif and WT as well as plotted acetylene reduction rates. Midpoint time represents the time to the inflection point of a logistic curve fit to the growth data (27), which highlights the extended growth lag in ancNif. Average growth parameter values are tabulated (five biological replicates per strain) ±1 SD. Bar plot shows mean acetylene reduction rates (three biological replicates per strain) and error bars represent ±1 SD. Asterisks indicate *p* < 0.05 (one-way ANOVA, post-hoc Tukey HSD). (A-C). Figures modified from Garcia et al. (20).

Strain ancNif was previously shown to exhibit a comparable diazotrophic growth rate to that of WT *A. vinelandii* albeit with a ∼14-hr growth lag (**Fig. 1B**) (20). Growth rates were assessed under molybdenum-replete conditions, under which the hybrid Mo-nitrogenase is expressed and the alternative V- and Fe-nitrogenases are repressed (8). Mo-nitrogenase activity in ancNif was found to be diminished (∼40% that of WT), as detected by decreased cellular reduction rates of the nitrogenase substrate, acetylene (**Fig. 1C**). This reduction in activity was previously confirmed by both nitrogenase immunodetection (**Fig. 1D**) and purified enzyme activity assays to be due to reduced nitrogenase activity rather than reduced protein concentration (20). Taken together, strain ancNif displays a measurable phenotypic defect while still retaining sufficient diazotrophic activity to sustain cell growth, making it a suitable target for transcriptional investigations into sequence-level nitrogenase perturbations.

### Gene expression patterns within the nitrogen fixation network are resilient to nitrogenase perturbations

We analyzed the transcriptome of the engineered ancNif strain sampled under mid-log, diazotrophic growth conditions by RNA-seq. Relative to WT strains cultured under the same conditions, we identified 405 genes (out of 5,051 mapped genes; ∼8%) that are significantly differentially expressed (adjusted *p*-value < 0.05), with 293 genes (∼6%) having increased transcript abundance in ancNif and 112 genes (∼2%) with decreased transcript abundance in ancNif (**Fig. 2, SI File 2**). Of those genes exceeding this significance threshold, 57 genes had at least a log_2_ fold-change of 2 in the ancNif strain compared to WT, either increased (54 genes) or decreased (3 genes) transcript abundance. We identified five clusters of gene response types in ancNif: three clusters among the 293 genes with increased transcript abundance in in ancNif and two clusters among the 112 genes with decreased transcript abundance in ancNif (**Fig. 2B, SI File 3**).

**Fig. 2.**
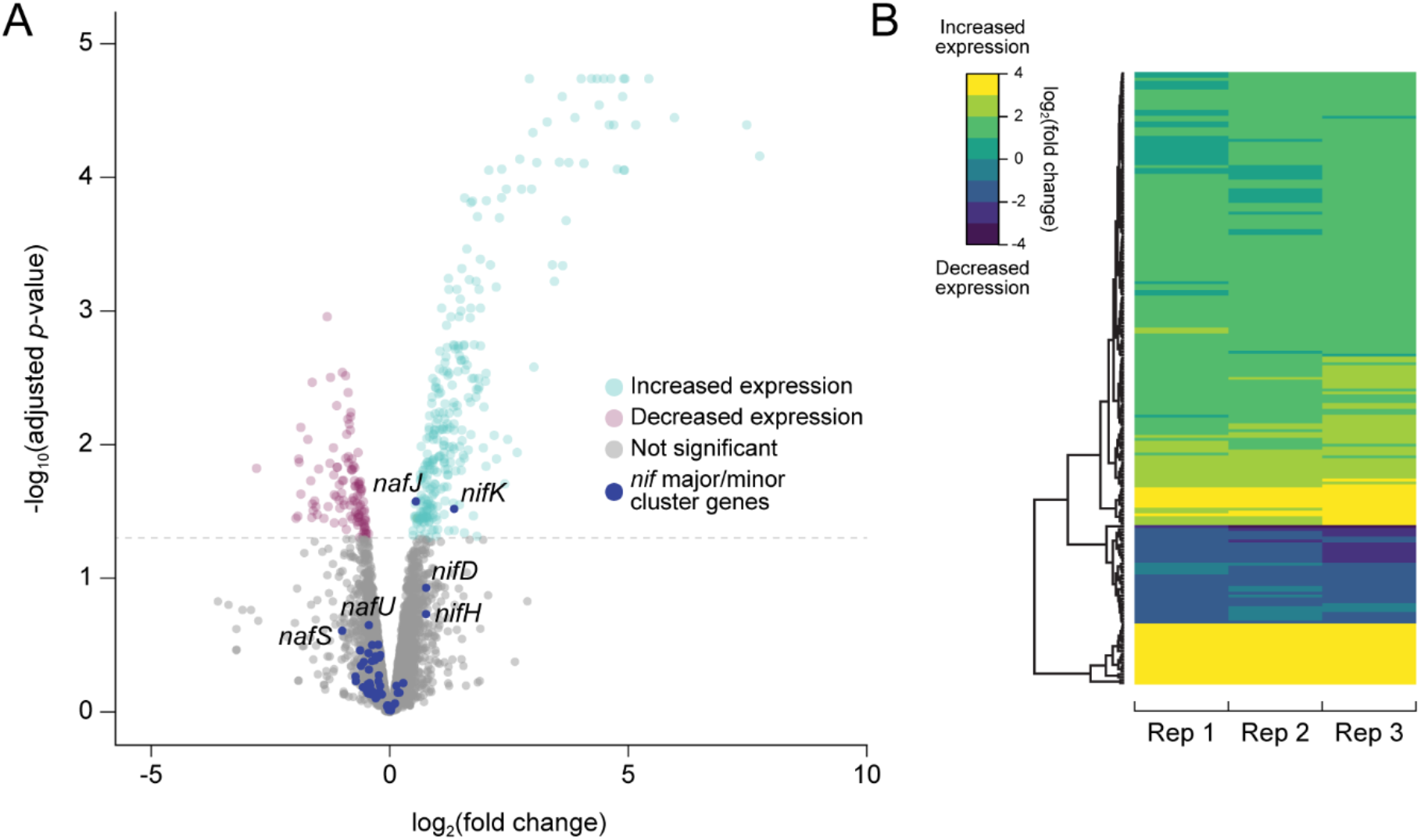
Global differential gene expression in strain ancNif relative to WT. (A) Volcano plot highlighting significantly differentially expressed genes in ancNif versus WT, defined by an adjusted *p* < 0.05. Data points corresponding to *nif* major or minor cluster genes are indicated in dark blue (see Fig. 3). (B) Clustering of gene expression patterns across three biological replicates of ancNif.

Because our genetic manipulation targeted the catalytic *nifD* gene, we investigated whether nitrogen-fixation-related genes were among those differentially expressed in the ancNif strain. We hypothesized that this manipulation would result in modifications to protein stoichiometries within the immediate nitrogen fixation network, accommodating a hybrid nitrogenase enzyme with reduced catalytic activity. In *A. vinelandii,* genes within the immediate network of the molybdenum-dependent, nitrogenase are arranged into two distinct clusters, the *nif* major (Avin_01360 to Avin_01720) and minor clusters (Avin_50900 to Avin_51060) (10, 28) (**Fig. 3**). The major cluster includes the *nifHDK* catalytic genes, and both clusters include other *nif* genes important for nitrogenase assembly, metallocluster biosynthesis, and regulation. Interspersed among *nif* cluster genes are *naf* genes (nitrogenase associated factors) that support assembly and biosynthesis, though many are not strictly required for diazotrophy and/or have unknown functions (10). The minor cluster contains *rnf* genes that form one of two respiratory complexes that direct electron flow to nitrogenase in *A. vinelandii* (the other is encoded by *fix* genes located elsewhere on the genome) (29). Predicted operons within the *nif* major and minor clusters mirror those previously reported (10, 30) (**SI File 4**). Finally, the dedicated genes for V- and Fe-nitrogenases in *A. vinelandii* are housed in *vnf* and *anf* clusters, respectively, though nitrogen fixation by these alternative nitrogenases has previously been shown to also be supported by expression of certain *nif* genes (8, 10, 26).

**Fig. 3.**
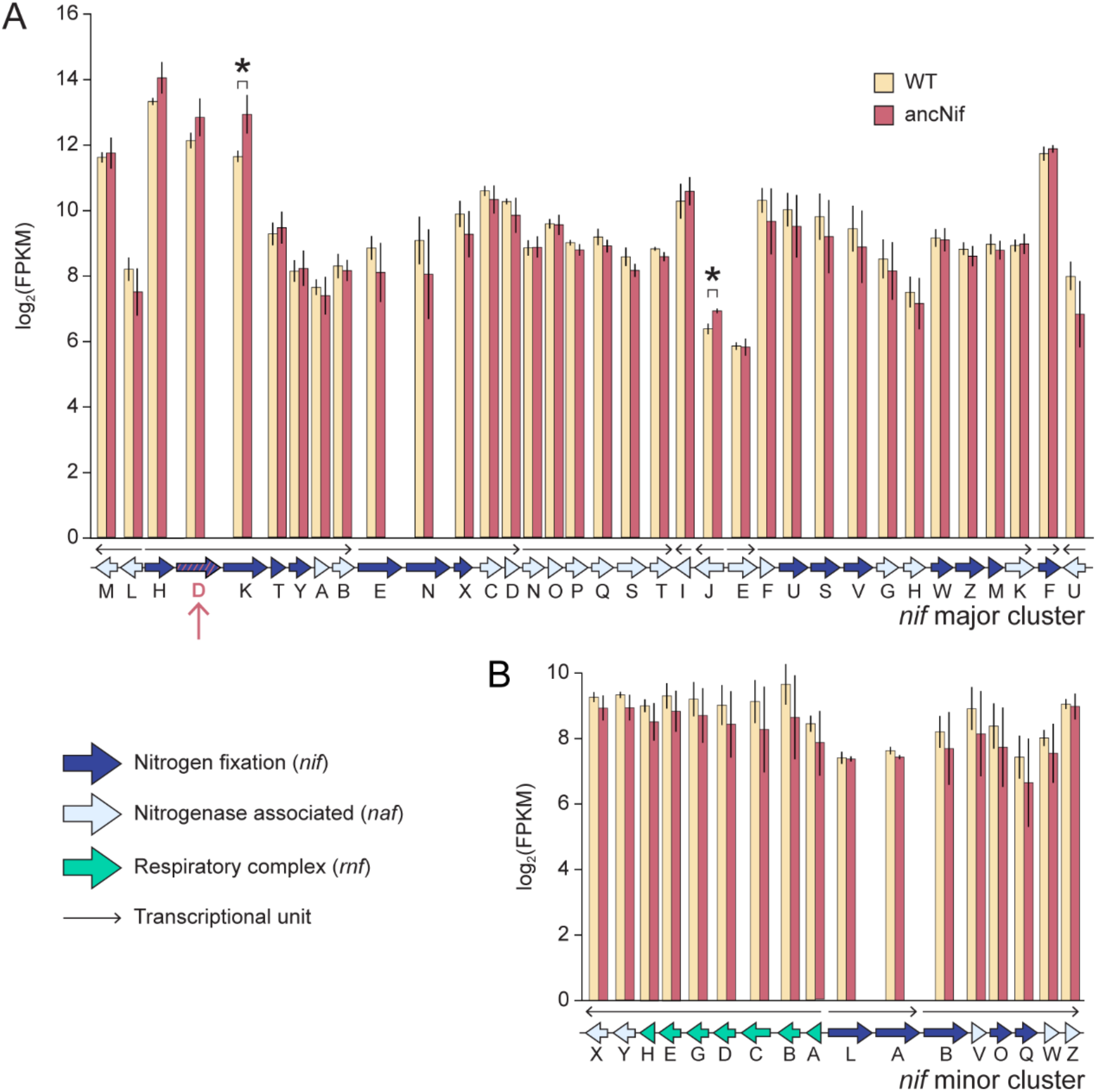
Transcription levels across the *nif* major and minor clusters of ancNif and WT *A. vinelandii* strains, expressed as fragments per kilobase per million mapped reads (FPKM). Bars represent mean values across three biological replicates per strain, and error bars indicate ± 1 SD. Asterisks indicate adjusted *p* < 0.05. Gene and transcriptional unit annotations from Del Campo et al. (10), the latter which mirrors operon predictions based on the transcriptional data presented here (**SI File 4**).

In both WT and ancNif strains, the catalytic nitrogenase genes, *nifHDK*, are the most highly expressed (log_2_(FPKM) ≈ 12-14, top 1% of all gene expression levels) relative to other genes within the major and minor *nif* clusters. In fact, in ancNif, *nifH* is the fourth most highly expressed gene in our dataset (*nifD* and *nifK* ranked within the top twelve), supporting previous findings that nitrogenase subunits constitute a significant percentage of total protein in nitrogen-fixing *A. vinelandii*. Little expression was observed for *vnf* and *anf genes* (e.g., ∼600-fold decreased expression of catalytic *vnfHDK* or *anfHDK* genes relative to *nifHDK*), which was expected since alternative nitrogen fixation in *A. vinelandii* is predicted to be repressed under the tested, molybdenum-replete, diazotrophic conditions (8).

Contrary to our hypothesis, we found that the expression of genes within the immediate nitrogen fixation network to be largely unaffected by replacement of an ancestral NifD variant in ancNif. Only two genes within the *nif* major and minor clusters showed significant increases in expression: the nitrogenase catalytic gene, *nifK* (Avin_01400; fold change ≈ 2.4), and an uncharacterized gene with an ABC transporter domain, *nafJ* (Avin_01580; fold change ≈ 1.5) (**Fig. 3, SI File 2**). Thus, relative stoichiometries of gene products across the *nif* clusters are expected to remain unchanged in ancNif. No genes within the *nif, vnf*, or *anf* clusters were found to have significantly lower expression in ancNif compared to WT. Together, these results suggest that nitrogen-fixation-related gene expression is remarkably robust to sequence-level perturbation to nitrogenase.

### Regulatory response is enriched in cellular functions outside the immediate nitrogen fixation network

We expanded our focus to analyze the regulatory response across cellular functions outside of the immediate nitrogen fixation network (i.e., outside of the major and minor *nif* clusters) that might otherwise be indirectly impacted by perturbation of the nitrogenase enzyme. We found several genes encoding proteins that are associated with pilus formation, molybdenum transport, electron transport, and central carbon metabolism to be among those with the largest expression changes in ancNif relative to WT (**Fig. 4**).

**Fig. 4.**
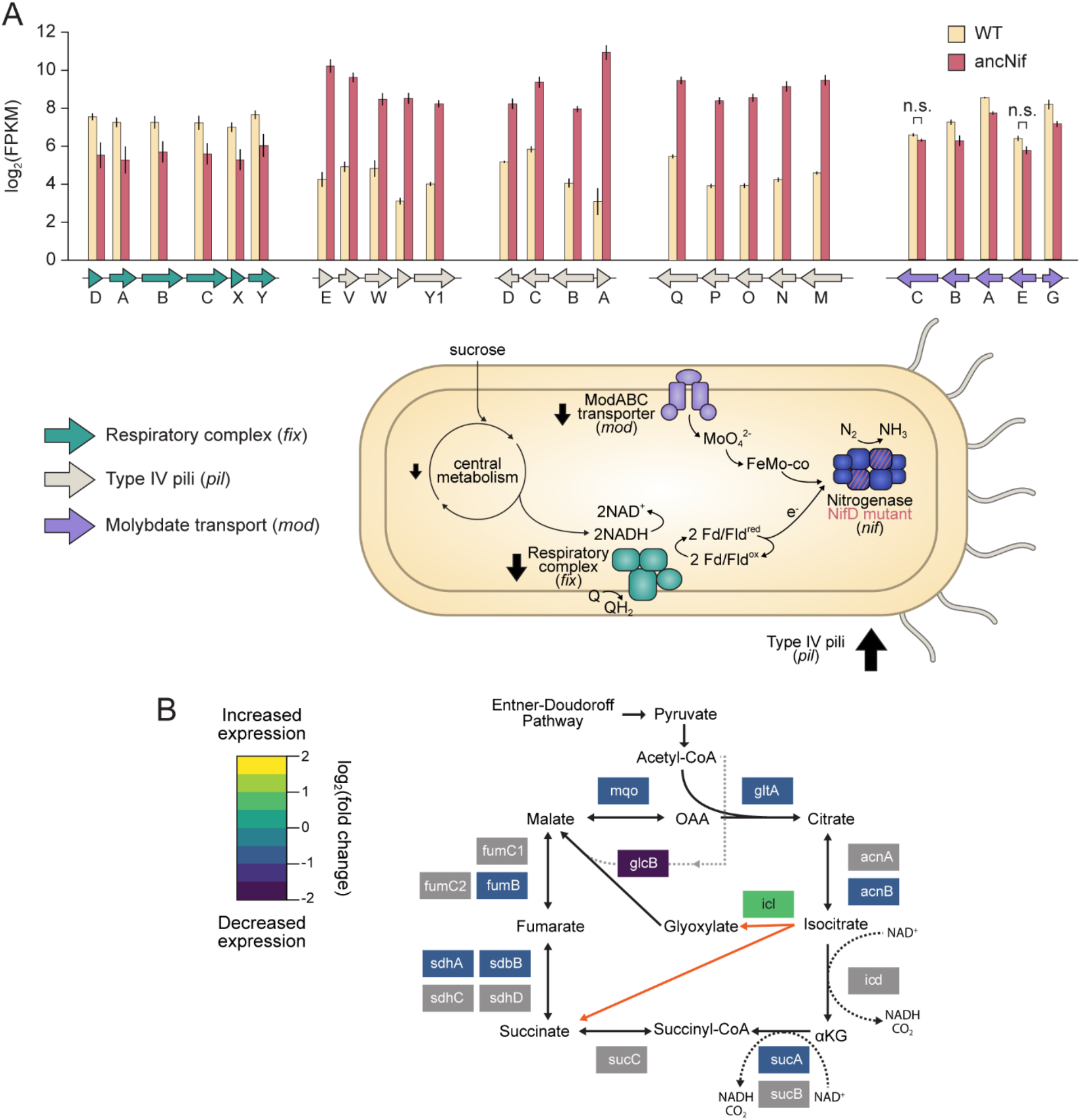
Differentially expressed genes encoding proteins involved in cellular functions external to the immediate nitrogen fixation network in ancNif. (A) Transcript levels across representative clusters related to respiration, motility, stress response, and molybdate transport. Bars represent mean values across three biological replicates per strain, and error bars indicate ± 1 SD. adjusted *p* < 0.05 for all genes except those labeled as not significant (“n.s.”). Schematic illustrates the relevance of cellular functions to nitrogen fixation in *A. vinelandii*. Arrows next to each cellular component signify associated gene expression changes relative to WT, with arrow thickness roughly corresponding to the magnitude of change and arrow direction indicating either increased expression in ancNif (arrow pointing upward) or decreased expression in ancNif (arrow pointing downward). (B) Transcription fold change levels mapped to the TCA cycle. Gene products colored in grey are not significantly differentially expressed in ancNif (adjusted *p* < 0.05). The glyoxylate shunt, mediated by the *icl* gene product, is highlighted in orange.

### Pilus formation

Of the 50 genes with the largest increases in expression in ancNif (∼1% of all mapped genes), 16 are related to Type IV pilus structure, assembly, and regulation, and have fold changes > 4 relative to WT. In fact, the gene with the largest increase in expression in ancNif (∼230-fold increase in expression relative to WT) is *pilA,* which codes for the major, filament-forming pilin protein (31). Other *pil* genes with increased expression are primarily located in three clusters (*pilEVWY1,* Avin_11830 to Avin_11870*; pilABCD*, Avin_12070 to Avin_12104; *pilMNOPQ*, Avin_45260 to Avin_45300; **Fig. 4A**). Across diverse bacteria, Type IV pili contribute to cell motility, sensing, attachment, aggregation, and DNA uptake (31). Further, large increases in Type IV pilus gene expression have previously been observed in the transition from non-diazotrophic to (Mo-dependent) diazotrophic conditions in *A. vinelandii* (8). The precise functional relationship between pilus formation and nitrogen fixation in *A. vinelandii* remains unclear. However, our finding that pilus gene expression is amplified even further in ancNif—exhibiting defects in its diazotrophic growth behavior relative to WT—might be indicative of resource limitation and an associated stress response. Indeed, increased Type IV pilus expression has previously been reported in bacteria under starvation conditions (32–34).

### Molybdenum transport

We observed that genes encoding proteins associated with molybdenum transport are expressed at lower levels (∼2-fold decrease) in ancNif relative to WT. *A. vinelandii* harbors a relatively complex pathway for molybdenum uptake and homeostasis that includes the transmembrane, high-affinity molybdenum ABC-type transporters (*mod*), as well as ATP-dependent molybdenum storage proteins (*mos*) (11, 35). In ancNif, *modABCGE* genes within the *mod1* cluster (Avin_50650 to Avin_50690; duplicates of some of these genes are also found within additional *mod2* and *mod3* clusters that are expressed at <10% the level of *mod1* in ancNif), which encode both the structural components of the transporter and its transcriptional repressor (*modE1*) (7), show lower expression levels (**Fig. 4**). Deletions of *modE1* and *mosAB* have each previously been shown to impair accumulation of intracellular molybdenum (35–37). Therefore, we expect that reduced expression of these genes in ancNif has a similar effect. Our results are consistent with a reduced cellular investment in accumulating high levels of intracellular molybdenum in ancNif that would otherwise support the molybdenum-dependent, *nif* nitrogenase system.

### Electron transport

Among the genes with the largest decrease in expression in ancNif relative to WT are those that encode the Fix respiratory complex, which, together with the Rnf complex, provides low potential electrons for nitrogen fixation (29, 38). Relative to WT, *fixABCX* genes (Avin_10520 to Avin_10560) in ancNif have ∼3 to 4-fold decreases in expression (**Fig. 4A**). The electron bifurcating Fix system transfers electrons from NADH to both quinone and flavodoxin/ferredoxin, the latter which donates electrons to nitrogenase (38). By contrast, Rnf, the expression of which is not significantly impacted in ancNif, couples reduction of flavodoxin/ferredoxin to the proton motive force. Previous deletion mutants have demonstrated that there is some degree of redundancy between the Rnf and Fix systems, but inactivation of both the *fix* and the *rnf* genes within the *nif* minor cluster is sufficient to abolish nitrogen fixation activity (38). Under varying oxygen conditions or expression of different nitrogenase isozymes, *A. vinelandii* cells display preferences for either Rnf or Fix as a means of optimizing electron flow for efficient nitrogen fixation. Fix is favored when oxygen is limiting and/or when additional energy is required for expression of alternative V- and Fe-nitrogenases (29). The decrease in Fix gene expression but relative maintenance of Rnf transcript levels in ancNif suggests both that these cells do not require the additional electron flux to nitrogenase that would have been generated by Fix and that these strains have a preference for the reducing power provided by Rnf.

### Central carbon metabolism

Several genes that encode enzymes within the *A. vinelandii* TCA cycl*e* exhibited decreased expression levels in ancNif compared to WT (∼2 to 3-fold decrease in expression; adjusted *p* < 0.05; **Fig. 4B, SI File 1 – Fig. S4**). We note, however, that expression of isocitrate lyase (*icl;* Avin_28420) was increased by ∼2-fold in ancNif. *icl* directs carbon toward the glyoxlate shunt and away from oxidative steps of the TCA cycle (39) that would otherwise generate reduced NADH for electron flow to nitrogenase. The expression changes within the TCA cycle in ancNif are opposite from what has previously been observed in mutant *A. vinelandii* strains that constitutively express nitrogenase. Constitutive nitrogenase expression leads to an increase in the expression of genes that encode enzymes in the TCA cycle, as well as decreased flux through the glyoxylate shunt (40). Whereas TCA cycle gene expression in the constitutively expressing *A. vinelandii* strain likely reflects greater cellular investment in generating reducing power for larger quantities of nitrogenase enzyme, the opposite outcome in ancNif instead implies a preference for reserving carbon via the gloxylate shunt at the expense of decreasing total electron flux toward nitrogen fixation.

### Reduced nitrogenase activity is compensated by allocation of fewer resources toward nitrogen fixation

Transcriptomic profiling of the mutant *A. vinelandii* strain ancNif harboring an ancestral NifD variant suggests a global pattern in which engineered cells reduce resources allocated for molybdenum-dependent nitrogen fixation in response to the diminished activity of the hybrid nitrogenase complex. Specifically, we find evidence that gene expression changes in the engineered microbe restrict molybdenum transport and storage, as well as electron flux by both downregulating Fix and upregulating the glyoxylate shunt that bypasses oxidative steps of the TCA cycle (**Fig. 4**). Though the specific relationship between increased expression of Type IV pilus genes and nitrogen fixation remains unclear, these patterns imply cellular investment in strategies to respond to resource limitation. Interestingly, once overcoming an extended growth lag, diazotrophic growth rates of the engineered microbe are indistinguishable from WT. Our results indicate that the lag may be an outcome of these metabolic rearrangements, which nevertheless eventually succeed in recovering typical growth rates despite reduced total nitrogenase activity (**Fig. 1**).

Under standard diazotrophic conditions, growth rate of *A. vinelandii* is limited by nitrogen fixation rate (29). Thus, the mutations in the ancestral, catalytic *nifD* gene of ancNif that slow nitrogenase activity are likely responsible for the initial lag in growth and would be expected to result in a wasteful excess of inputs to nitrogen fixation, including trace metals and reducing equivalents. The global expression patterns that we observe in ancNif demonstrate that this bottleneck in net nitrogen fixation rate is not—and perhaps cannot— be loosened by increasing the pool of assembled and active nitrogenase, as transcript levels of genes within the *nif* clusters remain relatively constant compared to WT (**Fig. 3**). A possible explanation for this is that WT *A. vinelandii* already expresses nitrogenase proteins at high levels (13), limiting the possible regulatory space that can be accessed for *nif* genes. Further, adjustments to stoichiometries of nitrogen-fixation-related genes may be similarly inaccessible, as *A. vinelandii,* like other well-studied diazotrophs, likely already expresses well-optimized ratios of *nif* gene products (16). Rather, we find that fitness reduction in engineered ancNif is most significantly managed by restricting cellular processes adjacent to nitrogen fixation, that are likely subject to more dynamic regulatory optimization.

## CONCLUSION

This transcriptomic dataset showcases the ability of *A. vinelandii* to readily incorporate phylogenetically inferred, ancient nitrogenase protein variants and respond dynamically to reduced nitrogen fixation activity in order to maintain diazotrophic growth rates. The regulatory response in an *A. vinelandii* strain engineered with an ancestral *nifD* sequence is enriched in genes external to the nitrogen fixation network, including those related to molybdenum processing and electron transport, as expression of *nif* genes is evidently resilient to nitrogenase perturbations and their physiological outcomes. This work provides a novel perspective on the challenge associated with engineering of *nif* gene expression for improvements to nitrogen fixation. Importantly, transcriptional patterns in the hybrid nitrogenase engineered strain highlight adjacent, upstream cellular processes as potentially more effective engineering targets for fine-tuning nitrogen fixation metabolism in bacteria, particularly in the interest of optimizing compatibilities between host models and nitrogenase protein variants.

## MATERIALS AND METHODS

### Ancestral sequence construction

Phylogenetic inference and reconstruction of the ancestral NifD variant sequence in strain ancNif was performed previously by Garcia et al. (20). Briefly, a representative nitrogenase protein sequence dataset was curated following an initial BLASTp (41) search of the NCBI non-redundant protein database. A concatenated NifHDK alignment was generated by MAFFT v7.450 (42) and phylogenetic inference and ancestral sequence reconstruction were performed by RAxML v8.2.10 (43) under the LG+G+F evolutionary model. Though this version of the RAxML software does not perform full marginal ancestral sequence reconstruction, the ancestral NifD protein sequence was previously not found to differ substantially when generated by the marginal reconstruction algorithm implemented in PAML (see Garcia et al. (20) for additional discussion).

### Nitrogenase structure prediction

The three-dimensional structure of the hybrid NifDK heterotetramer was predicted by Colabfold (44) (https://github.com/YoshitakaMo/localcolabfold), which combines the AlphaFold2 structure prediction method (45) with the MMSeq2 method for homology detection (46). Colabfold was run with standard options (3 recycles and AMBER all-atom optimization in GPU). Protein structures were visualized by ChimeraX (47).

### *A. vinelandii* genome engineering

**Table S1** contains information for strains and plasmids used in this study. *A. vinelandii* strain ancNif was previously constructed through markerless replacement of the WT *A. vinelandii* DJ *nifD* gene with an ancestral variant (20). Codon-optimized ancestral *nifD* and homologous flanking sequences were synthesized and cloned into a pUC19 vector (Twist Biosciences), generating plasmid pAG14. pAG14 was used to transform *A. vinelandii* DJ2278 (Δ*nifD::*KanR; generously provided by Dennis Dean, Virginia Tech), a non-diazotrophic mutant strain. The *nifD* variant was integrated into the genome via homologous reciprocal recombination, replacing the kanamycin resistance cassette within the *nifHDK* cluster. Transformants were screened for loss of kanamycin resistance and recovery of a diazotrophic phenotype. Desired insertion of the ancestral *nifD* gene was confirmed by Sanger sequencing of the *nifHDK* cluster (see **Table S2** for a list of primers).

### *A. vinelandii* growth and acetylene reduction

Phenotypic characterization of *A. vinelandii* strains in this study was performed previously (20), following Carruthers et al. (48). *A. vinelandii* WT, ancNif, and DJ2278 strains were cultured under Mo-replete, diazotrophic conditions. Cells were inoculated into flasks containing Burk’s medium lacking a fixed nitrogen source and grown at 30 °C and 300 rpm (five biological replicates per strain). Optical density at 600 nm (OD600) was monitored over the growth period. Growth parameters were estimated by the Growthcurver R package (27). To quantify cellular acetylene reduction rates, the strains described above were cultured to an OD600 ≈ 0.5 (three biological replicates per strain). Flasks were subsequently sealed and 25 mL of headspace was replaced by acetylene gas. Cultures were shaken at 30 °C and agitated at 300 rpm for 60 min. Headspace gas was sampled every 15 minutes over this period and analyzed on a Nexis GC-2030 gas chromatograph (Shimadzu). Acetylene reduction rates were normalized to total protein, quantified by the Quick Start Bradford Protein Assay kit (Bio-Rad).

### RNA library preparation and sequencing

Seed cultures of *A. vinelandii* WT and ancNif strains were grown in nitrogen-supplemented Burk’s medium (13 mM ammonium acetate) for 24 h at 30 °C and 300 rpm. Seed cultures were then inoculated into nitrogen-free Burk’s medium and grown diazotrophically to an OD600 of ∼0.7 (mid-log), immediately followed by RNA extraction with the RNeasy Mini kit (Qiagen) following manufacturer instructions. RNA extracts were assessed on a Nanodrop 2000c (Thermofisher Scientific) and confirmed to contain >2.5 µg of RNA with a purity of A260/280 = 1.8-2.2; A260/230 > 1.8. RNA extracts were stored at -80 °C. RNA library preparation and paired-end sequencing (2 x 150 bp read length; Illumina NovaSeq 6000) was performed by Novogene.

### RNA sequencing bioinformatic analysis

Raw FASTQ reads were trimmed using Trimmomatic version 0.3 (49) (default settings except for HEADCROP = 5, LEADING = 3, TRAILING = 3, SLIDINGWINDOW = 3:30, MINLEN = 36). Trimmed reads were aligned to the *A. vinelandii* DJ genome sequence (GenBank accession CP001157.1) using bwa-mem v0.7.17 (version 0.7.17-h5bf99c6_8) (50) with default parameters. Alignment files were further processed with Picard-tools v2.26.10 (https://broadinstitute.github.io/picard/) (CleanSAM and AddOrRepleaceReadGroups commands) and samtools v1.2 (51) (sort and index commands). Paired aligned reads were mapped to gene locations using HTSeq v0.6.0 (52) with default parameters. The R package edgeR v3.30.3 (53) was used to identify significantly differentially expressed genes from pairwise analyses, using Benjamini and Hochberg (54) adjusted *p*-value (FDR) < 0.05 as a significance threshold. Raw sequencing reads were normalized using the fragments per kilobase per million mapped reads method (FPKM). Volcano plots were constructed with ggplot2 (55). For significantly differentially expressed genes, clustering analysis of expression patterns was performed with DP_GP_cluster v0.1 (56). Operons were predicted using Operon-mapper (https://biocomputo.ibt.unam.mx/operon_mapper/, accessed 2022-10-19) (57) with default settings. The R package Pathview (58) was used to map fold changes of differentially expressed genes (FDR-adjusted *p* < 0.05) involved in the *A. vinelandii* TCA cycle to the KEGG (59) pathway map (avn00020). KEGG Gene Set Enrichment Analysis (GSEA) was performed using the R package clusterProfiler v.4.6.2 (60). The number of minimum required genes was set to 5 and the significant enrichment cutoff set to an FDR-adjusted *p* < 0.05.

### Data availability

Supplementary information for the present study is available in the following files:

SI_File_1.pdf – contains Figs. S1-S4 and Tables S1-S2
SI_File_2.xlsx – edgeR and FPKM analysis summary
SI_File_3.xlsx – expression clustering analysis summary
SI_File_4.xlsx – operon prediction summary

Phylogenetic tree from which the ancestral NifD was inferred and the predicted structure of the hybrid ancestral nitrogenase complex are available at https://github.com/kacarlab/ancNif_rna-seq_2023. Raw and processed RNA sequencing data are available at the NCBI GEO database (Accession pending).

## ACKNOWLEDGEMENTS

We thank Dennis Dean for providing A. vinelandii strains DJ and DJ2278; Jean-Michel Ané, April MacIntyre, and Junko Maeda for instrumentation support for acetylene reduction assays and helpful discussions; and Erica Majumder and Tim Donohue for valuable feedback. This research was supported by the National Aeronautics and Space Administration (NASA) Interdisciplinary Consortium for Astrobiology Research: Metal Utilization and Selection Across Eons, MUSE ICAR (#80NSSC21K0592), Great Lakes Bioenergy Research Center, U.S. Department of Energy, Office of Science, Office of Biological and Environmental Research (#DE-SC0018409), the National Science Foundation (#2228495) and the NASA Postdoctoral Program (MS).

## REFERENCES

1. Seefeldt LC, Yang ZY, Lukoyanov DA, Harris DF, Dean DR, Raugei S, Hoffman BM. 2020. Reduction of Substrates by Nitrogenases. Chem Rev 120:5082–5106. doi:10.1021/acs.chemrev.9b00556.

2. Rucker HR, Kacar B. 2023. Enigmatic evolution of microbial nitrogen fixation: insights from Earth’s past. Trends Microbiol doi:10.1016/j.tim.2023.03.011. doi:10.1016/j.tim.2023.03.011.

3. Fowler D, Coyle M, Skiba U, Sutton MA, Cape JN, Reis S, Sheppard LJ, Jenkins A, Grizzetti B, Galloway JN, Vitousek P, Leach A, Bouwman AF, Butterbach-Bahl K, Dentener F, Stevenson D, Amann M, Voss M. 2013. The global nitrogen cycle in the twenty-first century. Philos Trans R Soc Lond B Biol Sci 368:20130164. doi:10.1098/rstb.2013.0164.

4. Erisman JW, Galloway J, Seitzinger S, Bleeker A, Butterbach-Bahl K. 2011. Reactive nitrogen in the environment and its effect on climate change. Current Opinion in Environmental Sustainability 3:281–290. doi:10.1016/j.cosust.2011.08.012.

5. Vicente EJ, Dean DR. 2017. Keeping the nitrogen-fixation dream alive. Proc Natl Acad Sci U S A 114:3009–3011. doi:10.1073/pnas.1701560114.

6. Bennett EM, Murray JW, Isalan M. 2023. Engineering Nitrogenases for synthetic nitrogen fixation: From pathway engineering to directed evolution. BioDesign Research doi:10.34133/bdr.0005. doi:10.34133/bdr.0005.

7. Demtroder L, Narberhaus F, Masepohl B. 2019. Coordinated regulation of nitrogen fixation and molybdate transport by molybdenum. Mol Microbiol 111:17–30. doi:10.1111/mmi.14152.

8. Hamilton TL, Ludwig M, Dixon R, Boyd ES, Dos Santos PC, Setubal JC, Bryant DA, Dean DR, Peters JW. 2011. Transcriptional profiling of nitrogen fixation in Azotobacter vinelandii. J Bacteriol 193:4477–86. doi:10.1128/JB.05099-11.

9. Dixon R, Kahn D. 2004. Genetic regulation of biological nitrogen fixation. Nat Rev Microbiol 2:621–31. doi:10.1038/nrmicro954.

10. Martin Del Campo JS, Rigsbee J, Bueno Batista M, Mus F, Rubio LM, Einsle O, Peters JW, Dixon R, Dean DR, Dos Santos PC. 2023. Overview of physiological, biochemical, and regulatory aspects of nitrogen fixation in Azotobacter vinelandii. Crit Rev Biochem Mol Biol doi:10.1080/10409238.2023.2181309:1-47. doi:10.1080/10409238.2023.2181309.

11. Buren S, Jimenez-Vicente E, Echavarri-Erasun C, Rubio LM. 2020. Biosynthesis of Nitrogenase Cofactors. Chem Rev 120:4921–4968. doi:10.1021/acs.chemrev.9b00489.

12. Robson RL, Postgate JR. 1980. Oxygen and Hydrogen in Biological Nitrogen Fixation. Annual Review of Microbiology 34:183–207. doi:10.1146/annurev.mi.34.100180.001151.

13. Dingler C, Kuhla J, Wassink H, Oelze J. 1988. Levels and activities of nitrogenase proteins in Azotobacter vinelandii grown at different dissolved oxygen concentrations. Journal of Bacteriology 170:2148–2152. doi:10.1128/jb.170.5.2148-2152.1988.

14. Zhang L, Liu X, Li X, Chen S. 2015. Expression of the N2 fixation gene operon of Paenibacillus sp. WLY78 under the control of the T7 promoter in Escherichia coli BL21. Biotechnol Lett 37:1999-2004. doi:10.1007/s10529-015-1874-5.

15. Taggart JC, Lalanne JB, Li GW. 2021. Quantitative Control for Stoichiometric Protein Synthesis. Annu Rev Microbiol 75:243–267. doi:10.1146/annurev-micro-041921-012646.

16. Smanski MJ, Bhatia S, Zhao D, Park Y, L BAW, Giannoukos G, Ciulla D, Busby M, Calderon J, Nicol R, Gordon DB, Densmore D, Voigt CA. 2014. Functional optimization of gene clusters by combinatorial design and assembly. Nat Biotechnol 32:1241–9. doi:10.1038/nbt.3063.

17. Stripp ST, Duffus BR, Fourmond V, Leger C, Leimkuhler S, Hirota S, Hu Y, Jasniewski A, Ogata H, Ribbe MW. 2022. Second and Outer Coordination Sphere Effects in Nitrogenase, Hydrogenase, Formate Dehydrogenase, and CO Dehydrogenase. Chem Rev 122:11900–11973. doi:10.1021/acs.chemrev.1c00914.

18. Barahona E, Jimenez-Vicente E, Rubio LM. 2016. Hydrogen overproducing nitrogenases obtained by random mutagenesis and high-throughput screening. Sci Rep 6:38291. doi:10.1038/srep38291.

19. Jiang X, Paya-Tormo L, Coroian D, Garcia-Rubio I, Castellanos-Rueda R, Eseverri A, Lopez-Torrejon G, Buren S, Rubio LM. 2021. Exploiting genetic diversity and gene synthesis to identify superior nitrogenase NifH protein variants to engineer N2-fixation in plants. Commun Biol 4:4. doi:10.1038/s42003-020-01536-6.

20. Garcia AK, Harris DF, Rivier AJ, Carruthers BM, Pinochet-Barros A, Seefeldt LC, Kacar B. 2023. Nitrogenase resurrection and the evolution of a singular enzymatic mechanism. eLife 12. doi:10.7554/eLife.85003.

21. Sloan DB, Warren JM, Williams AM, Kuster SA, Forsythe ES. 2022. Incompatibility and Interchangeability in Molecular Evolution. EcoEvoRxiv doi:10.32942/X27P46. doi:10.32942/X27P46.

22. Noar JD, Bruno-Barcena JM. 2018. *Azotobacter vinelandii*: the source of 100 years of discoveries and many more to come. Microbiology 164:421–436. doi:10.1099/mic.0.000643.

23. Einsle O, Rees DC. 2020. Structural Enzymology of Nitrogenase Enzymes. Chem Rev 120:4969–5004. doi:10.1021/acs.chemrev.0c00067.

24. Garcia AK, Kolaczkowski B, Kacar B. 2022. Reconstruction of Nitrogenase Predecessors Suggests Origin from Maturase-Like Proteins. Genome Biol Evol 14. doi:10.1093/gbe/evac031.

25. Garcia AK, McShea H, Kolaczkowski B, Kacar B. 2020. Reconstructing the evolutionary history of nitrogenases: Evidence for ancestral molybdenum-cofactor utilization. Geobiology 18:394–411. doi:10.1111/gbi.12381.

26. Mus F, Alleman AB, Pence N, Seefeldt LC, Peters JW. 2018. Exploring the alternatives of biological nitrogen fixation. Metallomics 10:523–538. doi:10.1039/c8mt00038g.

27. Sprouffske K, Wagner A. 2016. Growthcurver: an R package for obtaining interpretable metrics from microbial growth curves. BMC Bioinformatics 17:172. doi:10.1186/s12859-016-1016-7.

28. Setubal JC, dos Santos P, Goldman BS, Ertesvag H, Espin G, Rubio LM, Valla S, Almeida NF, Balasubramanian D, Cromes L, Curatti L, Du Z, Godsy E, Goodner B, Hellner-Burris K, Hernandez JA, Houmiel K, Imperial J, Kennedy C, Larson TJ, Latreille P, Ligon LS, Lu J, Maerk M, Miller NM, Norton S, O’Carroll IP, Paulsen I, Raulfs EC, Roemer R, Rosser J, Segura D, Slater S, Stricklin SL, Studholme DJ, Sun J, Viana CJ, Wallin E, Wang B, Wheeler C, Zhu H, Dean DR, Dixon R, Wood D. 2009. Genome sequence of Azotobacter vinelandii, an obligate aerobe specialized to support diverse anaerobic metabolic processes. J Bacteriol 191:4534–45. doi:10.1128/JB.00504-09.

29. Alleman AB, Garcia Costas A, Mus F, Peters JW, Glass JB. 2022. Rnf and Fix Have Specific Roles during Aerobic Nitrogen Fixation in Azotobacter vinelandii. Applied and Environmental Microbiology 88. doi:10.1128/aem.01049-22.

30. Jacobson MR, Brigle KE, Bennett LT, Setterquist RA, Wilson MS, Cash VL, Beynon J, Newton WE, Dean DR. 1989. Physical and genetic map of the major nif gene cluster from Azotobacter vinelandii. Journal of Bacteriology 171:1017–1027. doi:10.1128/jb.171.2.1017-1027.1989.

31. Craig L, Forest KT, Maier B. 2019. Type IV pili: dynamics, biophysics and functional consequences. Nat Rev Microbiol 17:429–440. doi:10.1038/s41579-019-0195-4.

32. Wang C, Chen W, Xia A, Zhang R, Huang Y, Yang S, Ni L, Jin F, Kivisaar M. 2019. Carbon Starvation Induces the Expression of PprB-Regulated Genes in Pseudomonas aeruginosa. Applied and Environmental Microbiology 85. doi:10.1128/aem.01705-19.

33. Bansal R, Helmus RA, Stanley BA, Zhu J, Liermann LJ, Brantley SL, Tien M. 2013. Survival During Long-Term Starvation: Global Proteomics Analysis of Geobacter sulfurreducens under Prolonged Electron-Acceptor Limitation. Journal of Proteome Research 12:4316–4326. doi:10.1021/pr400266m.

34. Steindler L, Schwalbach MS, Smith DP, Chan F, Giovannoni SJ. 2011. Energy Starved Candidatus Pelagibacter Ubique Substitutes Light-Mediated ATP Production for Endogenous Carbon Respiration. PLoS ONE 6. doi:10.1371/journal.pone.0019725.

35. Navarro-Rodriguez M, Buesa JM, Rubio LM. 2019. Genetic and Biochemical Analysis of the Azotobacter vinelandii Molybdenum Storage Protein. Front Microbiol 10:579. doi:10.3389/fmicb.2019.00579.

36. Premakumar R, Jacobitz S, Ricke SC, Bishop PE. 1996. Phenotypic characterization of a tungsten-tolerant mutant of Azotobacter vinelandii. Journal of Bacteriology 178:691–696. doi:10.1128/jb.178.3.691-696.1996.

37. Mouncey NJ, Mitchenall LA, Pau RN. 1995. Mutational analysis of genes of the mod locus involved in molybdenum transport, homeostasis, and processing in Azotobacter vinelandii. Journal of Bacteriology 177:5294–5302. doi:10.1128/jb.177.18.5294-5302.1995.

38. Ledbetter RN, Garcia Costas AM, Lubner CE, Mulder DW, Tokmina-Lukaszewska M, Artz JH, Patterson A, Magnuson TS, Jay ZJ, Duan HD, Miller J, Plunkett MH, Hoben JP, Barney BM, Carlson RP, Miller AF, Bothner B, King PW, Peters JW, Seefeldt LC. 2017. The Electron Bifurcating FixABCX Protein Complex from Azotobacter vinelandii: Generation of Low-Potential Reducing Equivalents for Nitrogenase Catalysis. Biochemistry 56:4177–4190. doi:10.1021/acs.biochem.7b00389.

39. Dolan SK, Welch M. 2018. The Glyoxylate Shunt, 60 Years On. Annu Rev Microbiol 72:309–330. doi:10.1146/annurev-micro-090817-062257.

40. Wu C, Herold RA, Knoshaug EP, Wang B, Xiong W, Laurens LML. 2019. Fluxomic Analysis Reveals Central Carbon Metabolism Adaptation for Diazotroph Azotobacter vinelandii Ammonium Excretion. Sci Rep 9:13209. doi:10.1038/s41598-019-49717-6.

41. Camacho C, Coulouris G, Avagyan V, Ma N, Papadopoulos J, Bealer K, Madden TL. 2009. BLAST+: architecture and applications. BMC Bioinformatics 10:421. doi:10.1186/1471-2105-10-421.

42. Katoh K, Standley DM. 2013. MAFFT multiple sequence alignment software version 7: improvements in performance and usability. Mol Biol Evol 30:772–80. doi:10.1093/molbev/mst010.

43. Stamatakis A. 2014. RAxML version 8: a tool for phylogenetic analysis and post-analysis of large phylogenies. Bioinformatics 30:1312–3. doi:10.1093/bioinformatics/btu033.

44. Mirdita M, Schütze K, Moriwaki Y, Heo L, Ovchinnikov S, Steinegger M. 2022. ColabFold: making protein folding accessible to all. Nature Methods 19:679–682. doi:10.1038/s41592-022-01488-1.

45. Senior AW, Evans R, Jumper J, Kirkpatrick J, Sifre L, Green T, Qin C, Zidek A, Nelson AWR, Bridgland A, Penedones H, Petersen S, Simonyan K, Crossan S, Kohli P, Jones DT, Silver D, Kavukcuoglu K, Hassabis D. 2020. Improved protein structure prediction using potentials from deep learning. Nature 577:706–710. doi:10.1038/s41586-019-1923-7.

46. Steinegger M, Söding J. 2017. MMseqs2 enables sensitive protein sequence searching for the analysis of massive data sets. Nature Biotechnology 35:1026-1028. doi:10.1038/nbt.3988.

47. Pettersen EF, Goddard TD, Huang CC, Meng EC, Couch GS, Croll TI, Morris JH, Ferrin TE. 2021. UCSF ChimeraX: Structure visualization for researchers, educators, and developers. Protein Sci 30:70–82. doi:10.1002/pro.3943.

48. Carruthers BM, Garcia AK, Rivier A, Kacar B. 2021. Automated Laboratory Growth Assessment and Maintenance of *Azotobacter vinelandii*. Curr Protoc Microbiol 1:e57. doi:10.1002/cpz1.57.

49. Bolger AM, Lohse M, Usadel B. 2014. Trimmomatic: a flexible trimmer for Illumina sequence data. Bioinformatics 30:2114–2120. doi:10.1093/bioinformatics/btu170.

50. Li H, Durbin R. 2009. Fast and accurate short read alignment with Burrows– Wheeler transform. Bioinformatics 25:1754–1760. doi:10.1093/bioinformatics/btp324.

51. Li H, Handsaker B, Wysoker A, Fennell T, Ruan J, Homer N, Marth G, Abecasis G, Durbin R. 2009. The Sequence Alignment/Map format and SAMtools. Bioinformatics 25:2078–2079. doi:10.1093/bioinformatics/btp352.

52. Anders S, Pyl PT, Huber W. 2015. HTSeq—a Python framework to work with high-throughput sequencing data. Bioinformatics 31:166–169. doi:10.1093/bioinformatics/btu638.

53. Robinson MD, McCarthy DJ, Smyth GK. 2010. edgeR: a Bioconductor package for differential expression analysis of digital gene expression data. Bioinformatics 26:139–140. doi:10.1093/bioinformatics/btp616.

54. Benjamini Y, Hochberg Y. 1995. Controlling the False Discovery Rate: A Practical and Powerful Approach to Multiple Testing. Journal of the Royal Statistical Society: Series B (Methodological) 57:289–300. doi:10.1111/j.2517-6161.1995.tb02031.x.

55. Wickham H. 2009. ggplot2 doi:10.1007/978-0-387-98141-3.

56. McDowell IC, Manandhar D, Vockley CM, Schmid AK, Reddy TE, Engelhardt BE. 2018. Clustering gene expression time series data using an infinite Gaussian process mixture model. PLoS Comput Biol 14:e1005896. doi:10.1371/journal.pcbi.1005896.

57. Taboada B, Estrada K, Ciria R, Merino E, Hancock J. 2018. Operon-mapper: a web server for precise operon identification in bacterial and archaeal genomes. Bioinformatics 34:4118–4120. doi:10.1093/bioinformatics/bty496.

58. Luo W, Brouwer C. 2013. Pathview: an R/Bioconductor package for pathway-based data integration and visualization. Bioinformatics 29:1830–1. doi:10.1093/bioinformatics/btt285.

59. Kanehisa M. 2000. KEGG: Kyoto Encyclopedia of Genes and Genomes. Nucleic Acids Research 28:27–30. doi:10.1093/nar/28.1.27.

60. Yu G, Wang L-G, Han Y, He Q-Y. 2012. clusterProfiler: an R Package for Comparing Biological Themes Among Gene Clusters. OMICS: A Journal of Integrative Biology 16:284–287. doi:10.1089/omi.2011.0118.

