## Supplementary material for "A hybrid nitrogenase with regulatory elasticity in *Azotobacter vinelandii*": SI File 1

### Supplementary Information File 1

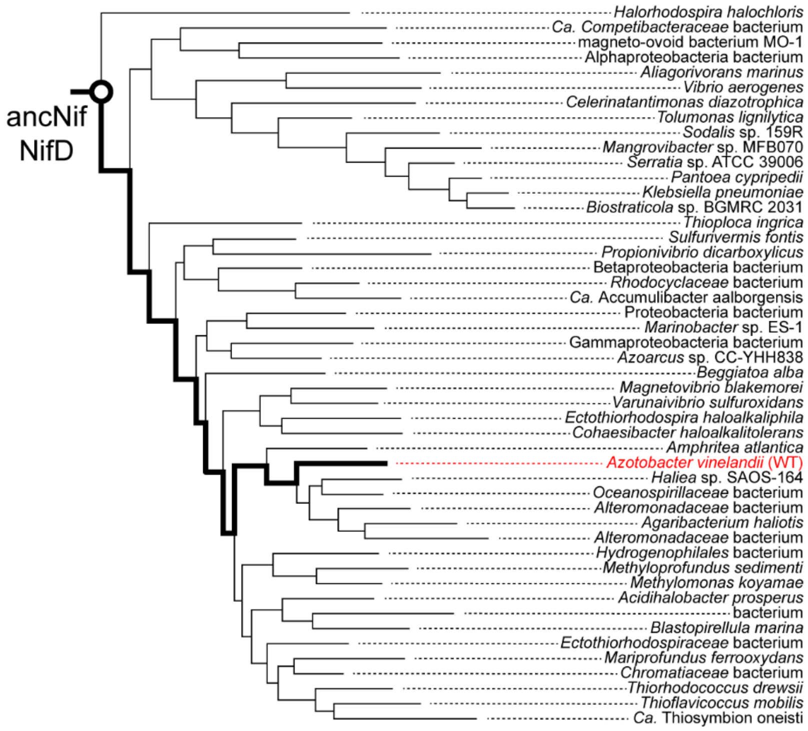


**Fig. S1.** Maximum likelihood phylogenetic subtree from which the ancestral ancNif NifD protein sequence was previously inferred. *A. vinelandii* lineage is highlighted with a bold line. Figure modified from Garcia et al. (1).

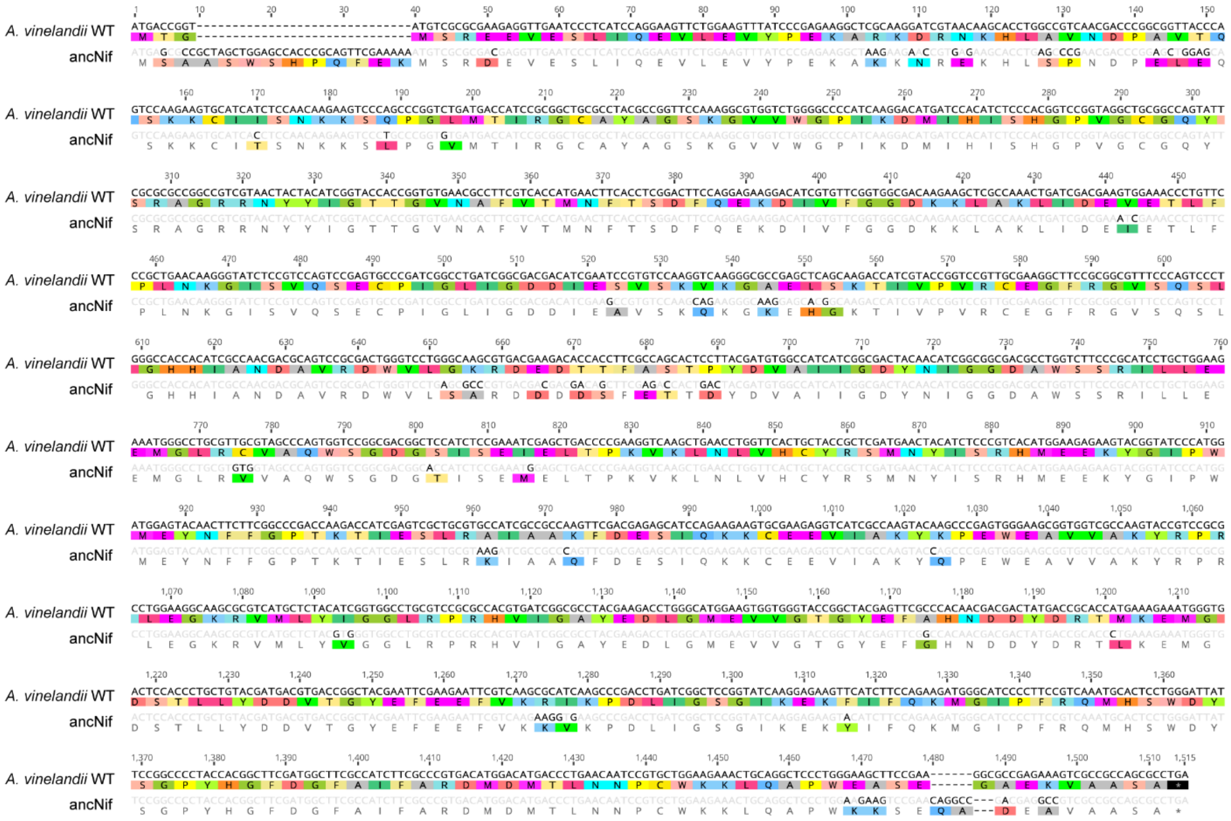


**Fig. S2.** Sequence alignment of WT and ancNif NifD proteins. Nucleotide and amino acid substitutions in the ancNif NifD protein relative to WT are highlighted.

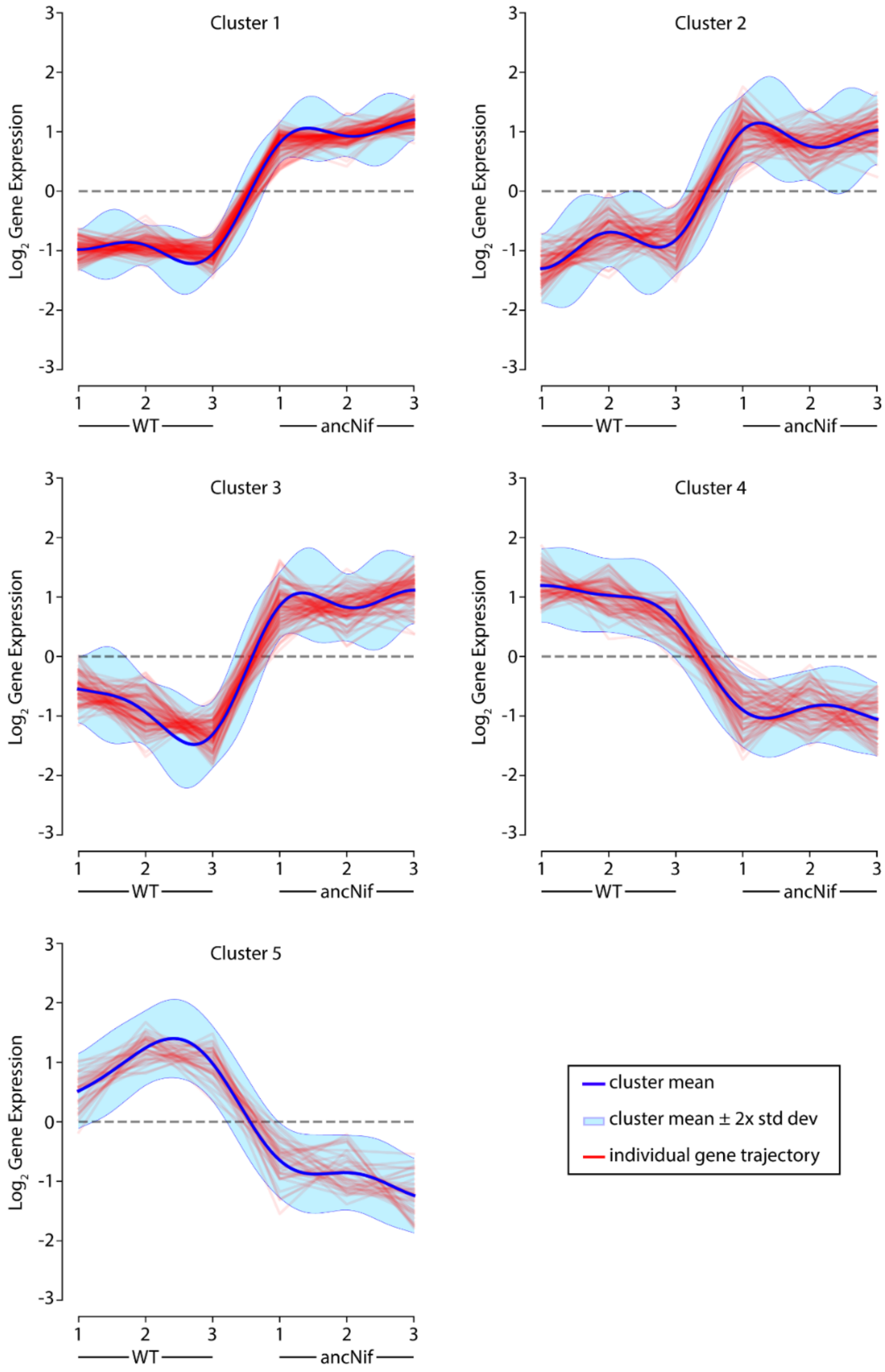


**Fig. S3.** Clustering analysis of significantly differentially expressed genes (FDR-adjusted *p*-value < 0.05 in ancNif relative to WT. Data from three biological replicates are represented for each strain.

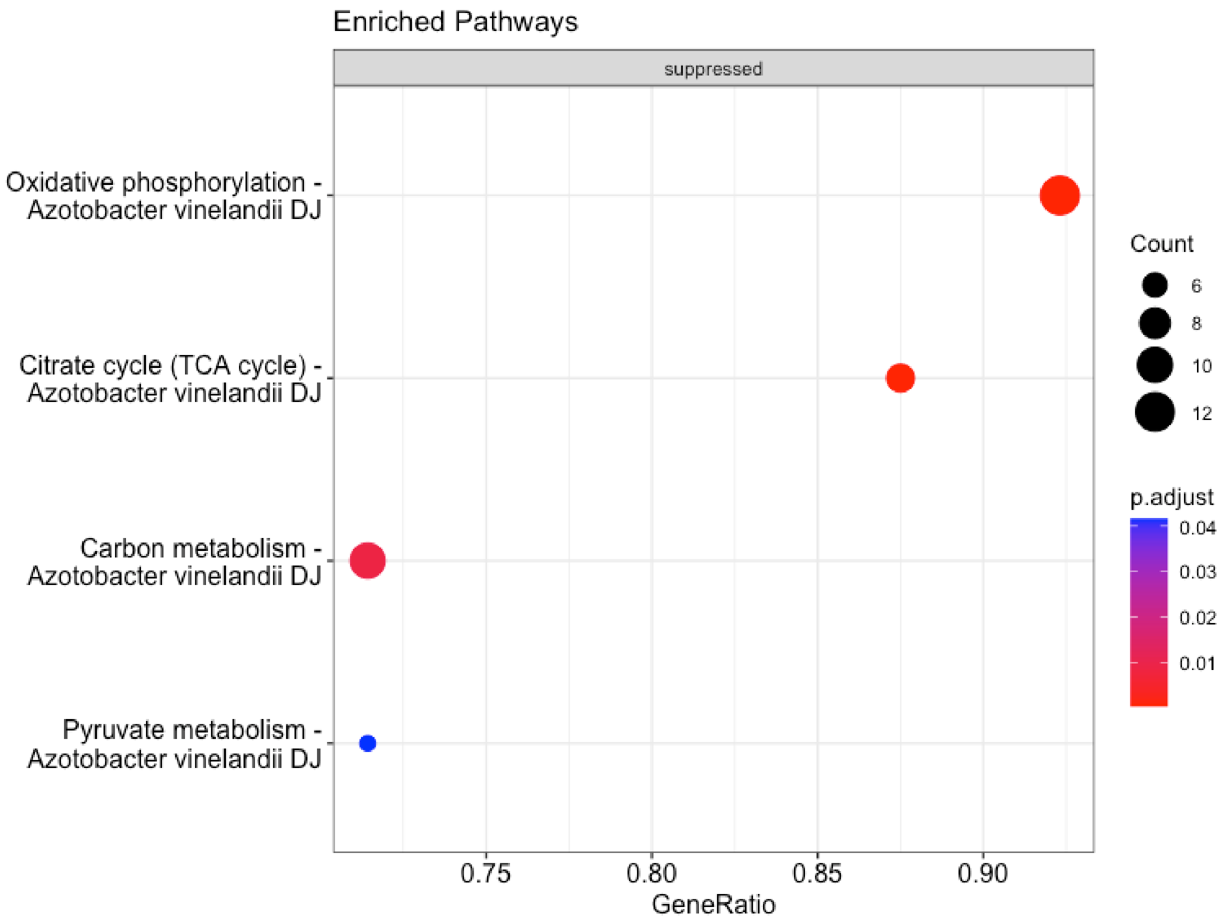


**Fig. S4**. KEGG pathways enriched for differential gene expression in ancNif relative to WT. KEGG pathway IDs: Oxidative phosphorylation (avn00190), Citrate cycle (avn0020), Carbon metabolism (avn01200), Pyruvate metabolism (avn00620).

**Table S1.** Strains and plasmids used in this study.

| **Name** | **Type** | **Description** | **Source** |
|---|---|---|---|
| DJ | Strain | Wild-type (WT); Nif+ | Dennis Dean, Virginia Tech |
| DJ2278 | Strain | Δ*nifD*::KanR; Nif- | Dennis Dean, Virginia Tech |
| ancNif* | Strain | Δ*nifD*::*nifD*$^{ancestor}$; RifR; Nif+; constructed by transforming DJ2278 with pAG14 | (1) |
| pAG14 | Plasmid | *nifD*$^{ancestor}$ + 400-bp *nifD* flanking homology regions, synthesized into XbaI/KpnI sites in pUC19 | (1) |

* Kaçar lab strain designation, “AK014”

**Table S2.** Primers used in this study.

| Primer | Sequence (5' to 3') | Description |
|---|---|---|
| 306_nifH_F | GCCGAACGTTCAAGTGGAAA | Forward primer, binds non-coding sequence upstream of *nifH; f*or PCR amplification of *nifHDK* and *nifH* sequencing |
| 307_nifH_R | AGAGCCAATCTGCCCTGTC | Reverse primer, binds non-coding sequence downstream of *nifH*; for *nifH* sequencing |
| 308_nifD_F | CACCCGTTACCCGCATATGA | Forward primer, binds non-coding sequence upstream of *nifD*; for *nifD* sequencing |
| 309_nifD_R | ACTCATCTGTGAACGGCGTT | Reverse primer, binds non-coding sequence downstream of *nifD*; for *nifD* sequencing |
| 310_nifK_F | GCTAACGCCGTTCACAGATG | Forward primer, binds non-coding sequence upstream of *nifK*; for *nifK* sequencing |
| 311_nifK_R | TCAGTTGGCCTTCGTCGTTG | Reverse primer, binds non-coding sequence downstream of *nifK*; for PCR amplification of *nifHDK* and *nifK* sequencing |

**REFERENCES**

1. Garcia AK, Harris DF, Rivier AJ, Carruthers BM, Pinochet-Barros A, Seefeldt LC, Kacar B. 2023. Nitrogenase resurrection and the evolution of a singular enzymatic mechanism. eLife 12. doi:10.7554/eLife.85003.
